# Widespread occurrence of ampicillin-susceptible *Enterococcus faecium* and *Enterococcus lactis* clinical isolates with low MICs to cephalosporins from Spain and Portugal

**DOI:** 10.64898/2026.08.28.747786

**Authors:** Miquel Sánchez-Osuna, Inmaculada Gómez-Sánchez, Juan C. Vázquez-Ucha, Ana C. Almeida-Santos, Paula Bierge, David Velasco, Judith Guitart-Matas, Silvia Capilla, Cristina Garcia-de-la-Mària, Salud Rodríguez-Pallares, Arianna Rodríguez-Coello, Antónia Read, Mariana Romanholo, Ana R. Freitas, Luisa Peixe, Oriol Gasch, Germán Bou, Carla Novais, Oscar Q. Pich

## Abstract

Reduced cephalosporin resistance in *Enterococcus faecium* has traditionally been reported in laboratory mutants and, more recently, in a single clinical ampicillin-susceptible (AmpS) isolate. Herein, we investigated whether this phenotype is widespread by analysing 95 clinical enterococcal isolates (78 AmpS and 17 ampicillin resistant [AmpR]) collected from three hospitals in Spain and Portugal (2009-2025). Low ceftriaxone MICs (≤4 mg/L) were detected in 19/51 (37.3%) AmpS *E. faecium* and 7/27 (25.9%) *E. lactis* but in none of the AmpR isolates. Low ceftriaxone MICs were associated with older patient age in both species and with prior ampicillin therapy in *E. faecium*, but not with other clinical or epidemiological variables. Ceftaroline MICs were consistently low among AmpS isolates, while ceftriaxone and cefotaxime showed greater variability. Low-MIC isolates were distributed across multiple clonal lineages and hospitals and did not share a distinctive resistance or virulence gene profile. PBP5 phylogeny and variation at the *psr*-*pbp5* region separated AmpS from AmpR *E. faecium* but did not explain variability in ceftriaxone MICs. Five AmpS isolates with reduced ceftriaxone MICs carried chromosomal deletions that included the *psr*-*pbp5* region and genes with diverse cellular functions. Variation in other candidate resistance genes (*pbpA*, *ponA*, *pbpF*, *croRS*, *stpA*/*stk* and *murAA*) did not consistently explain the MIC differences. These results reveal unexpected heterogeneity in intrinsic cephalosporin resistance in clinical *E. faecium* and *E. lactis* and suggest that additional genetic or regulatory mechanisms underlie reduced susceptibility.

## INTRODUCTION

*Enterococcus faecium* is recognized as one of the leading bacterial pathogens associated with healthcare-associated infections worldwide [1]. Its population has been classically structured into two major clades: clade A, comprising most hospital-associated strains, and clade B, primarily associated with the community strains. Recent studies indicate that clade B isolates actually correspond to *Enterococcus lactis* [2], which causes fewer human infections than *E. faecium*, and are currently misidentified as *E. faecium* in hospitals worldwide [3].

Although aminopenicillins remain a cornerstone of therapy for enterococcal infections, resistance is widespread among clinical *E. faecium* isolates but uncommon in other species, such as *E. lactis* [4]. High-level ampicillin resistance has been primarily associated with enhanced production of penicillin-binding protein 5 (PBP5) or expression of PBP5 variants with lower affinity for β-lactams [5,6]. In Europe, more than 75% of invasive *E. faecium* isolates are resistant to ampicillin [7], consistent with global trends reported in a recent large-scale analysis, which found ampicillin resistance in approximately 59% of clinical *E. faecium* strains worldwide, with prevalence increasing from 51.2% in 2000-2019 to 72.3% in 2020-2022 [8].

Beyond the widespread acquisition of ampicillin resistance, *Enterococcus* also displays intrinsic resistance to cephalosporins, rendering these antibiotics ineffective as monotherapy [9,10]. Although the molecular basis of cephalosporin resistance is not fully understood, several studies in *E. faecium* and *Enterococcus faecalis* have implicated genes encoding several PBPs such as *pbp5* [11], *pbpA* [12], *pbpF* and *ponA* [11], as well as *murAA*, involved in peptidoglycan synthesis [13], and the cell wall stress-responsive two-component systems *croRS* [14] and *stpA/stk* [15], all supported by data obtained from laboratory-generated mutants.

Our group has recently described few clinical ampicillin-susceptible *E. faecium* clinical isolates with unexpectedly low cephalosporin MICs originating from a single hospital epidemiological setting [16]. Now, we hypothesized that reduced cephalosporin MICs are not restricted to isolated clinical cases but represent a broader phenotype among ampicillin-susceptible *E. faecium*. Because *E. lactis* is the closest known relative of *E. faecium* [2], we also explored whether this phenotype extends to this recently recognized species. To test these hypotheses, we investigated the distribution and genomic diversity of ampicillin-susceptible *E. faecium* and *E. lactis* isolates with unusually low cephalosporin MICs across three hospitals in Spain and Portugal using antimicrobial susceptibility testing and whole-genome sequencing (WGS).

## MATERIAL AND METHODS

### Strains and clinical data

A total of 51 ampicillin-susceptible *E. faecium* and 27 *E. lactis* isolated between 2014 and 2025 were included in this study. Specifically, 22 *E. faecium* and 6 *E. lactis* isolates were obtained from Parc Taulí University Hospital (PTUH; Sabadell, Spain), 17 *E. faecium* and 4 *E. lactis* isolates from Complexo Hospitalario Universitario A Coruña (CHUAC; A Coruña, Spain) and 12 *E. faecium* and 17 *E. lactis* from Hospital Pedro Hispano (HPH; Porto, Portugal). Patient demographics and comorbidities, together with infection characteristics including source of isolation, co-infections with other bacteria and antibiotic therapy were recorded when available. Ampicillin-resistant *E. faecium* strains (n = 17) isolated between 2009 and 2025 were randomly selected for comparison, comprising 4 from PTUH, 1 from CHUAC and 12 from HPH.

### Antibiotic susceptibility testing

Susceptibility to ampicillin and ceftriaxone was assessed in all isolates using Etest strips (bioMérieux) according to EUCAST guidelines [17]. Ceftaroline and cefotaxime were also assessed in all PTUH isolates, which were representative of multiple genetic backgrounds and collection periods. In the absence of established clinical breakpoints for cephalosporins in enterococci, cephalosporin MICs below the ampicillin susceptibility breakpoint (≤ 4 mg/L) were here considered low MICs.

### Whole-genome sequencing

Isolates were cultured on Columbia agar plates supplemented with 5% sheep blood (bioMérieux) or Brain Heart Infusion (BHI) agar plates (Thermo Fisher Scientific) and incubated at 37°C. Genomic DNA was extracted using the DNeasy Blood & Tissue Kit (Qiagen) after a pre-treatment with 5 mg/mL lysozyme (Sigma-Aldrich) at 37°C for 30 minutes. DNA quality was assessed using a NanoDrop device (Thermo Fisher Scientific) and a Qubit fluorometer (Thermo Fisher Scientific).

Libraries for sequencing were prepared with the Nextera XT DNA Sample Preparation Kit (Illumina). WGS was performed using paired-end sequencing on an Illumina HiSeq 2500 and NovaSeq 600 sequencers. Raw sequencing read quality was assessed using FastQC v0.12.1 (https://github.com/s-andrews/FastQC). Reads were pre-processed and filtered with TrimGalore v0.6.6 (https://github.com/FelixKrueger/TrimGalore), followed by *de novo* assembly using shovill v1.1.0 (https://github.com/tseemann/shovill).

Nanopore sequencing (Oxford Nanopore Technologies, Oxford, UK) was performed according to the manufacturer’s instructions. Libraries were prepared using the ligation sequencing kit (LSK114) and native barcoding kit (NBD114-96) and sequenced on a PromethION P2 integrated system with an R10.4.1 flow cell. Basecalling and demultiplexing were performed using Dorado v7.9.8 (https://github.com/nanoporetech/dorado) in super-accurate mode. Reads shorter than 1 kb or with a quality score below Q10 were excluded. Sequencing adapters were removed prior to downstream analyses. Assemblies were generated using Unicycler v0.5.0 [18].

The resulting assemblies were evaluated for quality using CheckM2 v1.1.0 [19] and only those with estimated completeness ≥ 95% and contamination ≤ 5% were retained for further analyses. Genome dereplication was performed using dRep v2.2.3 with a secondary ANI threshold of 99.95% threshold [20]. Discrimination between *E. faecium* and *E. lactis* was assessed using FastANI v1.34 against the NCBI reference genomes *E. faecium* SRR24 and *E. lactis* CX 2-6_2, assigning each genome based on the best average nucleotide identity (ANI) hit, with a ≥ 95% ANI threshold for species assignment [21]. Genome annotation was performed with prokka 1.14.6 [22]. Multilocus sequence typing (MLST) was performed using the MLST v2.23.0 software (https://github.com/tseemann/mlst), based on both the classical [23] and the recently published MLST scheme for *E. faecium* [24]. MLST was not performed for *E. lactis* because no MLST scheme is currently available for this species. Novel sequence types (STs) were submitted to PubMLST for future reference. Putative acquired antibiotic resistance genes (ARG) and virulence factors (VF) were predicted on the assembled scaffolds with ABRicate v1.0.1 (https://github.com/tseemann/abricate) using the in-built Resfinder database [25] and the *E. faecium* and *E. lactis* database for VirulenceFinder [26] and otherwise default parameters. Prediction of mutations conferring antimicrobial resistance was conducted using ResFinder v4.5.0 against the *E. faecium* database [25].

### Comparative genomic analysis

To identify genes previously associated with low cephalosporin MICs, a comparative genomics analysis was performed using all ampicillin-susceptible *E. faecium* genomes included in this study. The genes analyzed included *pbp5* (E6A31_RS06825), *pbpA* (E6A31_RS03475), *ponA* (E6A31_RS06205), *pbpF* (E6A31_RS12065), *croR* (E6A31_RS13520), *croS* (E6A31_RS13525), *stpA* (E6A31_RS12560), *stk* (E6A31_RS12555) and *murAA* (E6A31_RS09060) [11–13,15,27]. Homologs were identified using reciprocal BLASTP v2.16.0+ searches [28], using the corresponding *E. faecium* SRR24 protein sequences as queries. Searches were performed using a conservative E-value threshold of <1e–20 and a query coverage ≥ 75%.

Given that truncations of *psr* have been reported [29], homologs of the *psr* gene were identified using BLASTN v2.16.0+ searches [28] with the full-length *psr* gene from *E. lactis* COM15 [CVT45_RS06345] as the query. Hits with ≥ 80% sequence identity and ≥ 30% query coverage were retained for further analysis. Synteny and conservation of the *psr* genetic environment were analyzed using clinker v0.0.32 [30].

### Phylogenetic inference

Core-genome phylogenetic analysis was performed based on single nucleotide polymorphisms (SNPs) called by snippy (https://github.com/tseemann/snippy) against the *E. faecium* SRR24 reference genome [GCF_009734005]. A ML tree was constructed with IQ-TREE 2 v2.0.7 [31] using 1000 bootstrap replicates and TVM+F+I+G4 as the substitution model, as deduced from ModelFinder [32].

Multiple sequence alignments of the protein products of gene homologs previously associated with cephalosporin resistance and of DNA regions containing *psr* and *pbp5* were generated with CLUSTALW 2.1 [33] using gap opening penalties of 5, 10 and 25. These alignments were integrated with local pairwise alignments generated by LALIGN using T-COFFEE 13.41.0.28bdc39 [34]. The resulting alignments were trimmed with Gblocks 0.91b [35] using the “half gap” setting. Maximum-likelihood (ML) phylogenetic trees were inferred with IQ-TREE 2 v2.0.7 [31], with substitution models selected using ModelFinder [32] and 1,000 bootstrap replicates. *Enterococcus hirae* FDAARGOS_1123 PBP5 protein sequence was used as the root, following a previously published PBP5 phylogeny [36].

Tree visualization and annotation were performed with iTOL v7 [37].

### Statistical analysis

Continuous variables were compared using the Mann-Whitney U test (MWU) and categorical variables were analyzed with Fisher’s exact test (FT). Correlations between MIC values were assessed using the Pearson correlation coefficient (PCC). A *p*-value < 0.05 was considered statistically significant. All statistical analyses were performed using custom Python scripts.

### Data availability

Whole-genome sequencing data generated in this study have been deposited in the NCBI database with accession number PRJNA1513992. Additionally, some strains included in this project were previously sequenced and are available under different projects (PRJNA1132796, PRJNA1290041).

## RESULTS

### Low cephalosporin MICs in ampicillin-susceptible *E. faecium* and *E. lactis*

Low ceftriaxone MICs (≤ 4 mg/L) were observed in 19/51 (37.3%) *E. faecium* and 7/27 (25.9%) *E. lactis* isolates (Figure 1, Data S1). By hospital, low ceftriaxone MICs were observed in 6/17 *E. faecium* and 2/4 *E. lactis* isolates at CHUAC (38.1%); 4/12 and 3/17 at HPH (24.1%); and 9/22 and 2/6 at PTUH (39.3%). Ceftriaxone MIC_50_ values were 32.0 mg/L for ampicillin-susceptible *E. faecium* and 24.0 mg/L for *E. lactis*.

**Figure 1.**
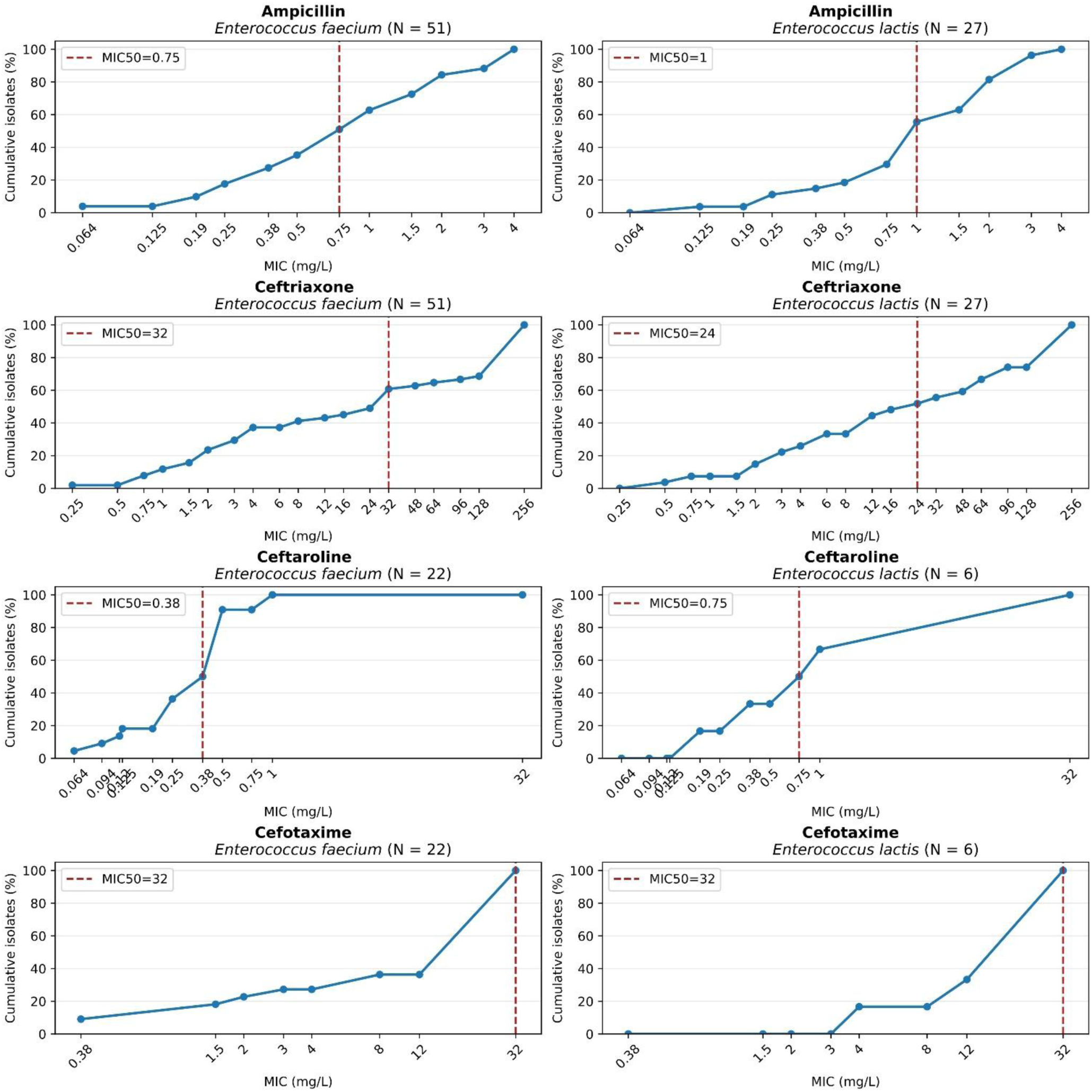
Cumulative MIC distributions of ampicillin-susceptible *E. faecium* and *E. lactis* for ampicillin, ceftriaxone, ceftaroline and cefotaxime. Curves represent the cumulative percentage of isolates inhibited at each MIC value (log₂ scale). Red dashed lines denote the MIC₅₀, defined as the lowest MIC inhibiting at least 50% of isolates within each species. Ceftaroline and cefotaxime MICs were determined only for isolates from PTUH. For plotting purposes, MIC values reported as >32 mg/L and >256 mg/L were considered as 32 mg/L and 256 mg/L, respectively.

Susceptibility testing of representative isolates revealed distinct patterns for ceftaroline and cefotaxime. All 22 ampicillin-susceptible *E. faecium* isolates tested and 4/6 *E. lactis* showed low ceftaroline MICs (Figure 1), whereas low cefotaxime MICs were observed in 6/22 and 1/6 isolates, respectively. Leveraging that these isolates were tested for ampicillin and three different cephalosporins (ceftriaxone, ceftaroline and cefotaxime), we assessed correlations between MIC values for these antibiotics. The near-perfect correlation between ampicillin and ceftaroline MICs (PCC = 0.98) contrasted with the only moderate correlation observed with the third-generation cephalosporins ceftriaxone and cefotaxime (PCC = 0.60) (Figure 2).

**Figure 2.**
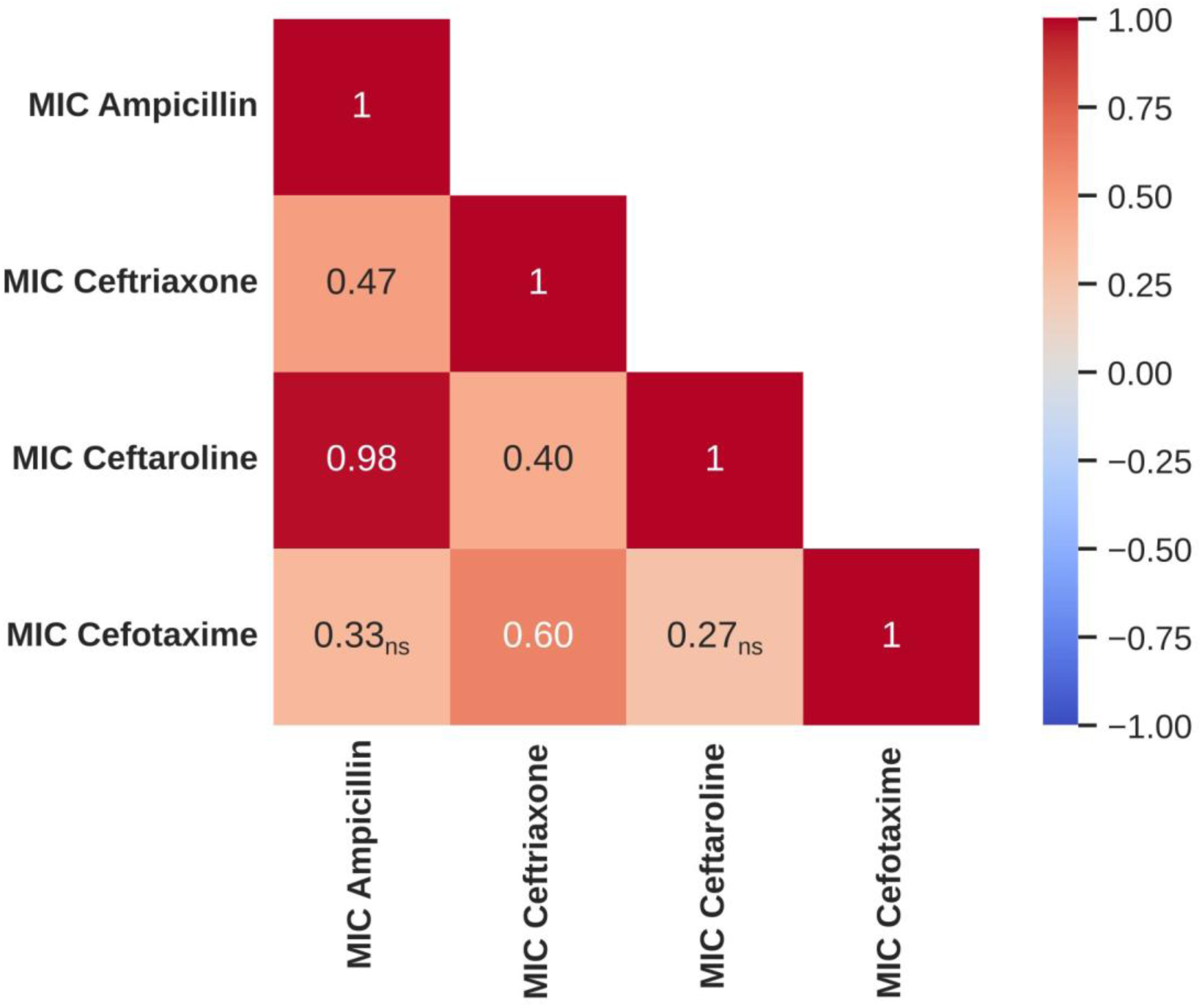
Pairwise Pearson correlations between MIC values for ampicillin, ceftriaxone, ceftaroline and cefotaxime in PTUH isolates. The color scale represents Pearson Correlation Coefficients (PCC), ranging from -1 (blue) to +1 (red). Non-significant correlations are indicated as “*ns*”.

To assess whether low cephalosporin MICs were restricted to ampicillin-susceptible enterococci, we included 17 ampicillin-resistant *E. faecium* to the study. All exhibited ceftriaxone MICs >256 mg/L. Again, representative isolates were also tested against ceftaroline and cefotaxime, showing uniformly the highest MIC values (>256 mg/L and >32 mg/L, respectively) (Data S2).

### Clinical and epidemiological data of ampicillin-susceptible isolates and features associated with low ceftriaxone MICs

The ampicillin-susceptible strains included in the present study were isolated between 2014 and 2025 (Data S1). Ceftriaxone MICs remained stable throughout the study period, with no correlation between MIC and year of isolation (PCC = −0.055). Of the corresponding patients, 50.0% (39/78) were female and 48.7% (38/78) were male. To one patient data was not available. The mean age was 70.9 ± 15.1 years. The most frequent clinical conditions were diabetes (17.9%, 14/78), arterial hypertension (12.8%, 10/78), dyslipidemia (11.5%, 9/78) and obesity (9.0%, 7/78). No comorbidities were identified in 7.7% (6/78) of patients and information was missing for 6.4% (5/78). The most common sample sources were blood (44.9%, 35/78), pus (19.2%, 15/78), bile (10.2%, 8/78), ascitic fluid (7.7%, 6/78) and wounds (5.1%, 4/78). A total of 50 strains (64.1%) were isolated from polymicrobial infections, with *Escherichia coli* being the most frequently co-isolated species (58.0%, 29/50). A total of 49 infections (62.8%) were treated with an antibiotic therapy combination, most frequently including piperacillin-tazobactam (35.9%, 28/78), meropenem (23.1%, 18/78), amoxicillin-clavulanate (20.5%, 16/78), linezolid (17.9%, 14/78), ceftriaxone (17.9%, 14/78), metronidazole (16.7%, 13/78), vancomycin (12.8%, 10/78), daptomycin (9.0%, 7/78) and ampicillin (9.0%, 7/78). Clinical and epidemiological data were not available for most of the ampicillin-resistant isolates (Data S2).

To explore potential clinical and epidemiological factors associated with ceftriaxone MICs, univariate analyses were performed among all ampicillin-susceptible isolates of *E. faecium* and *E. lactis* (Table S1). Patient age was significantly associated with ceftriaxone MIC group (MWU test, *p* < 0.05), with patients whose isolates belonged to the low-MIC group showing a higher median age than those whose isolates were in the high-MIC group (79.5 versus 67 years). This difference was also observed when analyzed separately by species, with median ages of 79 versus 66.5 years among *E. faecium* patients and 80 versus 70 years among *E. lactis* patients in the low- and high-MIC groups, respectively. Ampicillin treatment was also significantly less frequent in the high-MIC group than in the low-MIC group (2/52, 3.8%, versus 5/26, 19.2%; FT, *p* < 0.05). Species-specific analyses indicated that this association was driven exclusively by *E. faecium*. Specifically, four ampicillin-treated patients in the *E. faecium* subset belonged to the ceftriaxone low-MIC group, whereas none of the patients in the high-MIC group had received ampicillin. No significant associations were identified for the remaining clinical variables.

### Genomic and phylogenetic analysis of *E. faecium* and *E. lactis* isolates

According to the classical MLST scheme, ampicillin-susceptible and ampicillin-resistant *E. faecium* isolates were assigned to 38 and 11 different STs, respectively. In contrast, using the recently established MLST scheme, these isolates were classified into 50 and 14 STs (Data S1). Two *E. faecium* could not be assigned to a ST according to the new MLST scheme because they lacked the *mdlA* and *uvrA* genes, respectively.

Core-genome phylogenetic analysis (Figure 3) revealed a well-supported separation between *E. faecium* and *E. lactis* isolates. Isolates from the three hospitals were distributed throughout the tree, indicating the absence of clear phylogeographic structure. Ampicillin-resistant *E. faecium* tended to cluster together and were clearly separated from most ampicillin-susceptible isolates. In contrast, isolates exhibiting low ceftriaxone MICs were scattered throughout the phylogenetic tree, predominantly among ampicillin-susceptible isolates, indicating that this phenotype is not confined to specific clonal lineages, as supported by the MLST analysis.

**Figure 3.**
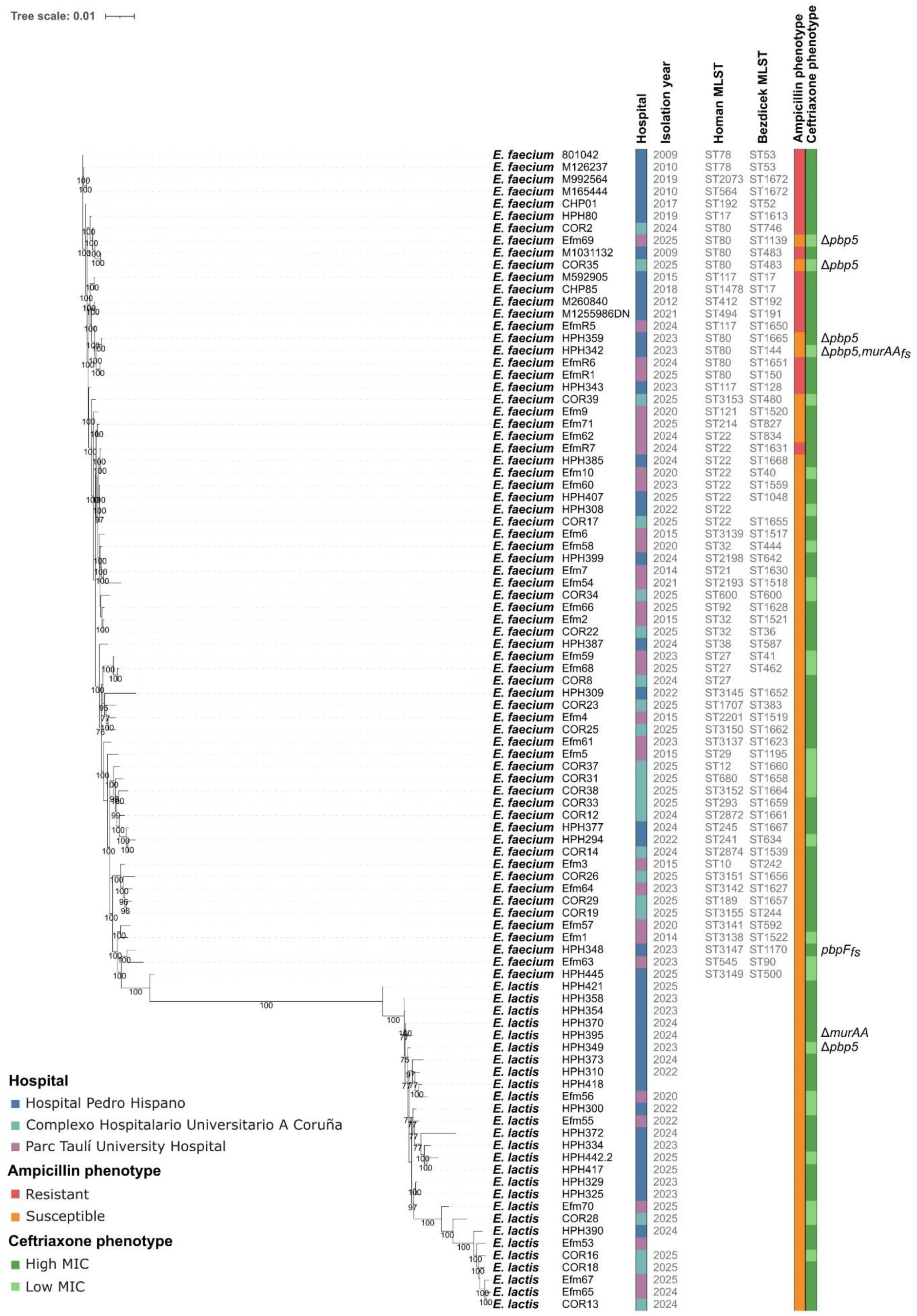
Core-genome phylogenetic tree of *E. faecium* and *E. lactis* isolates analyzed in this study. Branch support values represent the percentage of bootstrap replicates supporting each node and are shown only for branches with ≥ 75% support. Isolates are annotated according to hospital of origin, year of isolation, MLST, ampicillin susceptibility phenotype and ceftriaxone MIC category. Isolates lacking or carrying frameshift mutations in *pbp5*, *pbpF* or *murAA* are also indicated. Frameshift mutations are indicated as “*fs*”.

Among the ampicillin-susceptible *E. faecium* isolates, the most frequently detected acquired ARGs were *tet(M)* (45.1%, 23/51), *ant(6)-Ia* (27.5%, 14/51), *erm(B)* and *tet(L)* (25.5%, 13/51), *lsa(E)* (21.6%, 11/51), *aph(3′)-III* and *lnu(B)* (19.6%, 10/51) (Data S1). Despite being phenotypically ampicillin-susceptible, ResFinder identified PBP5 mutations predicted to confer ampicillin resistance in 47 isolates (92.1%). In addition, mutations in *gyrA* and *parC* potentially associated with resistance to ciprofloxacin were detected in 5 strains (6.4%). Regarding VFs, *lysM2-*Entfm and *lysM4-*Entfm were detected in all *E. faecium* isolates (100%, 51/51). Other highly prevalent VFs included *ccpA*-Entfm-hosp (94.1%, 48/51), *fms13*-Entfm (92.2%, 47/51), *bepA*-Entfm-com and *lysM3*-Entfm-hosp (90.2%, 46/51), *fnm*-Entfm-com, *gls20*-Entfm, *gls33*-Entfm, *glsB1*-Entfm and *glsB*-Entfm (88.2%, 45/51), *sagA*-Entfm-hosp (82.4%, 42/51), *empA*-Entfm-hosp (74.5%, 38/51), *empC*-Entfm-hosp and *fms17*-Entfm-com (72.5%, 37/51) or *lysM1*-Entfm-hosp (70.6%, 36/51). No significant differences in genetic features were observed between ampicillin-susceptible *E. faecium* isolates with low ceftriaxone MICs and those exhibiting high ceftriaxone MICs (Figure S1, Table S2). Similarly, no differences were observed in the mean number of ARGs, point mutations or VFs per isolate between the two groups (MWU, *p* > 0.05).

In *E. lactis*, the most frequently detected acquired ARGs were *tet(M)*, identified in two isolates (7.4%), and *tet(L)* and *vanNXY*, each detected in one isolate (3.7%) (Data S1). Among these three isolates, one exhibited a low ceftriaxone MIC, whereas the other two exhibited high ceftriaxone MICs. In contrast to *E. faecium*, PBP5 substitutions were less common in *E. lactis*, being detected in 13 isolates (48.1%). Regarding VFs, *ccpA*-Entls, *fnm*-Entls, *lysM2*-Entfm and *lysM4*-Entfm were detected in all *E. lactis* isolates (100%, 27/27). Other highly prevalent VFs included *acm*-Entfm-hosp and *gls33*-Entls (96.3%, 26/27), *gls20*-Entls and *glsB1*-Entls (88.9%, 24/27), and *glsB*-Entls and *sagA*-Entls (85.2%, 23/27). Additional VFs, including *bepA*-Entfm-com, *fms13*-Entls, *fms14*-Entfm-hosp, *fms16*-Entls and *fms19*-Entls, were detected in 70.4% (19/27) of isolates.

### Comparative genomic analysis of genes associated with cephalosporin resistance

With the exception of 5 isolates, most of *E. faecium* and *E. lactis* encoded a PBP5 homolog (Table S3). Phylogenetic analysis revealed a clear separation between ampicillin-susceptible and ampicillin-resistant isolates (Figure 4). Notably, the PBP5 substitutions M485A and E629V were present in all ampicillin-resistant isolates and absent from all susceptible isolates. Mean Needleman-Wunsch amino acid identity between PBP5 proteins from ampicillin-resistant and ampicillin-susceptible isolates was 97.0 ± 1.1% (Table S4). No association was observed between PBP5 clades and ceftriaxone MICs. Consistent with this, mean pairwise amino acid identity among PBP5 proteins was similar across ampicillin-susceptible isolates (98.4 ± 0.9%), isolates with low ceftriaxone MICs (98.4 ± 0.9%) and those with high MICs (98.5 ± 0.9%) (Table S4).

**Figure 4.**
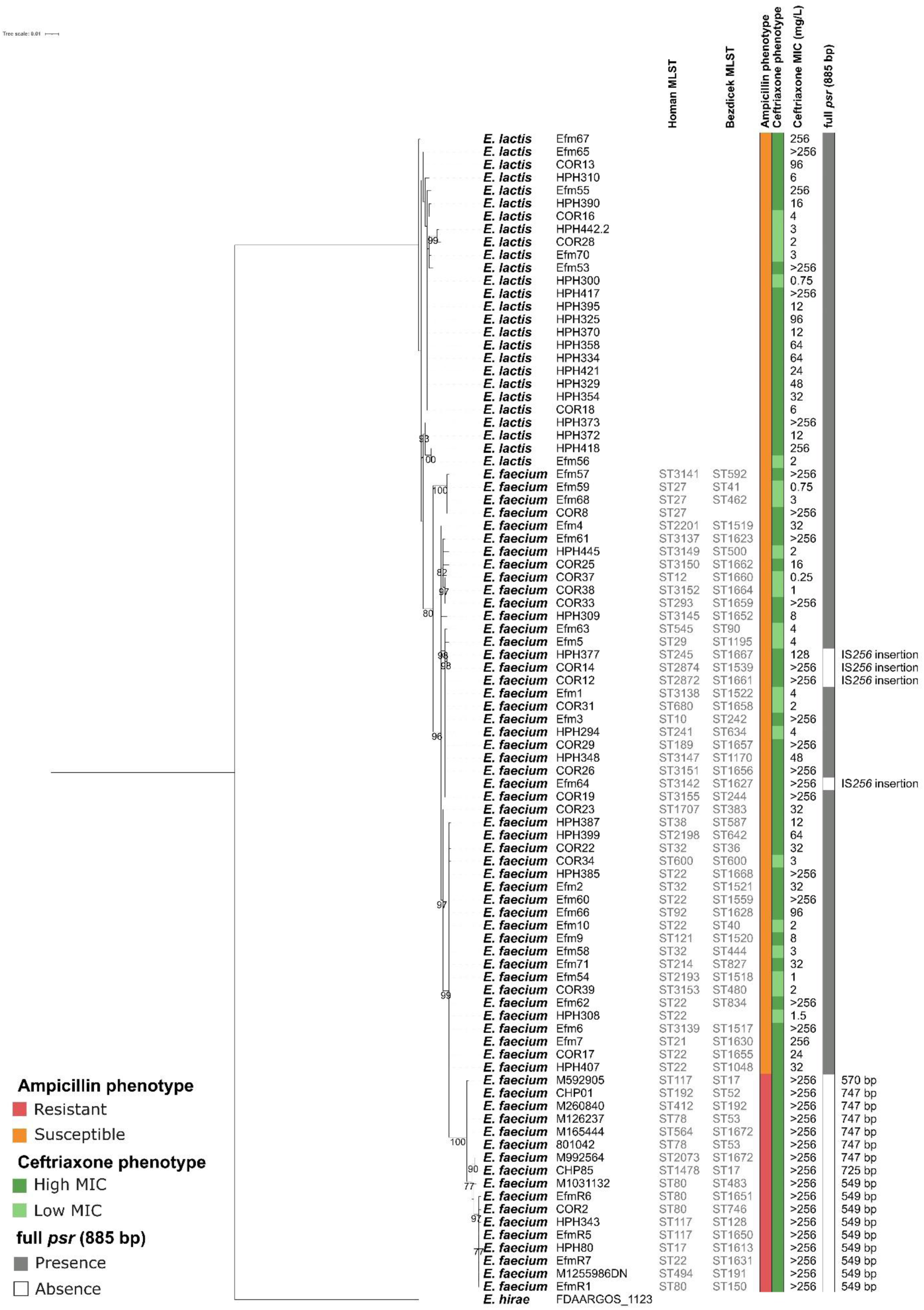
Rooted PBP5 phylogeny of *E. faecium* and *E. lactis* isolates analyzed in this study. Branch support values represent the percentage of bootstrap replicates supporting each node and are shown only for branches with ≥ 75% support. MLST, ampicillin susceptibility phenotype, ceftriaxone MIC and *psr* type are also indicated. *E. hirae* FDAARGOS_1123 PBP5 protein sequence was used as the outgroup.

Analysis of the *psr* locus, located upstream of *pbp5*, showed that 26 isolates lacked a full-length *psr* gene (Table S5). Specifically, 9 isolates carried a 549-bp *psr* allele, 6 carried a 747-bp allele, one carried a 570-bp allele and another carried a 725-bp allele. These truncated alleles were exclusively identified in ampicillin-resistant *E. faecium*. In contrast, 4 ampicillin-susceptible *E. faecium* isolates carried an IS*256* transposase insertion within *psr*. All these isolates lacking a full-length *psr* gene exhibited high ceftriaxone MICs (128 to >256 mg/L). Phylogenetic analysis of the *psr-pbp5* locus supported its association with ampicillin resistance (Figure S2). The 5 ampicillin-susceptible isolates lacking *pbp5* also lacked *psr*. Except for *E. lactis* HPH349, all *E. lactis* isolates encoded a full-length *psr* gene.

Further analysis of the ampicillin-susceptible isolates lacking both *psr* and *pbp5* revealed that this loss was associated with larger chromosomal deletions. Ampicillin-susceptible *E. faecium* isolates COR35, Efm69, HPH342 and HPH359 (ST80 according to the classical MLST scheme), carried chromosomal deletions of 32.3 to 57.2kb encompassing 39 to 60 genes when compared with the *E. faecium* SRR24 genome (Table S3). All deletions included the *ftsW-psr-pbp5* gene cluster, as well as multiple VFs (*glsB*-Entfm, *glsB1*-Entfm, *gls20*-Entfm, *gls33*-Entfm) and genes involved in phosphotransferase systems, carbohydrate metabolism, cell envelope biogenesis, membrane transport, DNA repair, iron-sulfur cluster biosynthesis and transcriptional regulation (Table S6). These strains clustered within the clade containing the ampicillin-resistant *E. faecium* isolates and were the only ampicillin-susceptible strains in this clade (Figure 3). A similar pattern was observed in *E. lactis* HPH349, which carried a 148.8kb deletion encompassing 133 genes relative to the *E. lactis* COM15 genome (Table S3). This deletion also included the *pbp5* operon, several VFs (*empA*-Entfm-hosp, *glsB*-Entls, *gls20*-Entls and *gls33*-Entls) and genes involved in cell envelope biogenesis, carbohydrate metabolism and transport, membrane transport, stress response, DNA repair, iron-sulfur cluster biosynthesis, protein secretion and folding, and transcriptional regulation (Table S6).

All isolates with *psr-pbp5* platform deletions were classified as low ceftriaxone MICs, except *E. faecium* HPH359, which showed a reduced MIC of 24 mg/L compared with the >256 mg/L typical of intrinsically resistant strains (Figure 3, Data S1). Despite differences in deletion size, all five deletions shared a core set of 29 genes, including the *psr-pbp5* locus and several VFs, including *glsB*, *gls20* and *gls33* genes, with upstream or downstream extensions depending on the isolate (Figure S3).

Other genes previously implicated in cephalosporin intrinsic resistance (*pbpA*, *ponA*, *croRS* and *stpA/stk*) were present in all ampicillin-susceptible isolates, including those with low cephalosporin MICs (Table S3). In contrast, *pbpF* carried a premature stop codon in the ampicillin-susceptible *E. faecium* HPH348 (ceftriaxone MIC of 48 mg/L), and *murAA* was deleted or frameshifted in HPH395 (MIC 12 mg/L) and HPH342 (MIC 1.5 mg/L), respectively. Phylogenetic analysis of the individual or concatenated protein sequences of these cephalosporin resistance-associated genes did not clearly resolve the ampicillin susceptibility phenotype or ceftriaxone MIC groups (Figure S4). Mean pairwise amino acid identity across the individual proteins ranged from 99.0% to 99.8%, reflecting the high sequence conservation of these loci, indicating a high degree of sequence conservation that may limit species-level resolution in the corresponding phylogenetic trees.

## DISCUSSION

Enterococci are traditionally considered intrinsically resistant to cephalosporins [9]. However, our multicentric data suggest that intrinsic high-level cephalosporin resistance is not universal among clinical ampicillin-susceptible *E. faecium* and *E. lactis*. Low cephalosporin MICs were identified in isolates recovered from three hospitals in Spain and Portugal, spanning multiple years and diverse phylogenetic backgrounds, indicating that this phenotype is neither geographically restricted nor confined to a particular clonal lineage. This suggests that reduced cephalosporin MICs represent a broader and previously underrecognized phenotype among clinical enterococci.

Most ampicillin-susceptible isolates exhibited low ceftaroline MICs, consistent with shared or partially overlapping determinants of susceptibility to these β-lactams. Similar observations have been reported previously, although only in a limited number of isolates [38], likely because cephalosporin susceptibility testing is not routinely performed for enterococci. Several *in vitro* studies have reported synergistic or enhanced activity of ampicillin plus ceftaroline against both *E. faecium* and *E. faecalis*, although results are isolate-dependent [38–40]. A similar synergistic relationship has not been consistently observed for ceftriaxone or cefotaxime, which aligns with ceftaroline’s distinct PBP-binding profile and relatively greater anti-enterococcal activity [41]. The strong correlation between ampicillin and ceftaroline susceptibility, together with the weaker correlations observed for ceftriaxone and cefotaxime, suggests that susceptibility to third-generation cephalosporins may involve additional or distinct determinants. Thus, ceftaroline susceptibility may be more predictable in ampicillin-susceptible isolates, whereas ceftriaxone and cefotaxime susceptibility appears more variable.

Reduced susceptibility to both ampicillin and cephalosporins has been largely attributed to alterations in the low-affinity penicillin-binding protein PBP5 [6,11,29]. Consistent with this, our phylogenetic analysis showed a clear association between PBP5 sequence variation and the ampicillin susceptibility phenotype. The PBP5 substitutions M485A and E629V, previously linked to ampicillin resistance, were found in all ampicillin-resistant isolates in our collection and were absent from susceptible strains [42]. However, because PBP5 amino acid identity was similar across low- and high-MIC groups in our dataset, sequence variation alone does not fully explain ceftriaxone susceptibility, supporting a multifactorial model.

Notably, we also identified isolates carrying large chromosomal deletions that include the *psr* and *pbp5* genes. These isolates were ampicillin-susceptible and had lower ceftriaxone MICs, suggesting that loss of the *psr*-*pbp5* region may increase susceptibility to both ampicillin and cephalosporins. Previous comparative genomics studies have shown that large chromosomal platforms containing *pbp5* and associated adaptive genes can be transferred between distinct *E. faecium* lineages, highlighting the mobility of this genomic region [43,44]. In this context, the identification of clinical isolates carrying large deletions encompassing *psr-pbp5* suggests that loss of this region may also occur naturally in clinical populations. Notably, the loss of this platform also resulted in the loss of multiple *gls* genes, which have been implicated in virulence and bile acid stress tolerance [45]. Thus, these deletions may have broader phenotypic consequences beyond their possible effects on β-lactam susceptibility, potentially affecting bacterial adaptation to the host environment.

A recent study reported that a full-length *psr* gene results in lower PBP5 expression and lower ampicillin MICs compared with shorter *psr* alleles, regardless of the *pbp5* allele present [29]. Consistent with this observation, isolates carrying full-length *psr* alleles in our collection were ampicillin-susceptible but showed substantial variability in ceftriaxone MICs. In contrast, *E. faecium* isolates with shorter *psr* alleles were ampicillin-resistant and also exhibited high ceftriaxone MICs. In such strains, we hypothesize that loss of a full-length *psr* allele leads to increased PBP5 expression and thereby contributes to elevated ceftriaxone MICs, consistent with our previous finding that higher PBP5 expression was associated with increased ceftriaxone MICs in single-point isolates from a naturally occurring *E. faecium* strain with a low baseline MIC [46]. Of note, isolates carrying IS*256* insertions that disrupted *psr* were ampicillin-susceptible despite exhibiting high ceftriaxone MICs, suggesting that ampicillin susceptibility in these isolates may not be explained solely by *psr* disruption and may instead involve additional mechanisms. In contrast, in isolates with complete loss of the *psr-pbp5* platform, the regulatory effect of *psr* on PBP5 would also be absent because *pbp5* is no longer present. Thus, the lower ampicillin and ceftriaxone MICs observed in these isolates may reflect the loss of PBP5 itself rather than a regulatory effect mediated by *psr*.

However, several isolates with a full-length *psr* gene also exhibited high ceftriaxone MICs, indicating that *psr* alone does not explain ceftriaxone susceptibility. In this context, we identified isolates carrying complete *psr* genes that showed reduced ceftriaxone MICs and harbored inactivating mutations in *pbpF* or *murAA*. PBPF is a class A PBP described as an essential partner of PBP5 in peptidoglycan polymerization; however, deletion of *pbpF* alone in *E. faecium* D344R did not reduce the ceftriaxone MIC [11]. By contrast, MurAA participates in the initial steps of peptidoglycan biosynthesis and its deletion has been associated with reduced cephalosporin susceptibility in *E. faecalis* [13]. These observations underscore that cephalosporin susceptibility is multifactorial and arises from the interplay of multiple genetic determinants.

Collectively, genes associated with cephalosporin resistance occurred in diverse combinations across our isolates, with some features shared between species. This diversity suggests that β-lactam resistance is highly strain-dependent and may result from different combinations of genetic determinants. Importantly, the findings presented in this study were obtained from clinical isolates, providing a distinct context from most studies focusing on reduced cephalosporin susceptibility in *Enterococcus*. Previous evidence linking point mutations, gene presence or absence and *psr-pbp5* variation to reduced cephalosporin MICs has largely been generated using genetically manipulated laboratory strains. In contrast, the genetic features identified in our study occurred naturally in clinical isolates. Although these observations do not necessarily establish causality, their occurrence in clinical populations supports their potential relevance to the variability in cephalosporin susceptibility observed in *E. faecium* and *E. lactis*. Expression and functional studies will be required to determine how these different genetic backgrounds and combinations of determinants contribute to the phenotype.

However, several limitations should be considered when interpreting our findings. First, the isolate collection was derived from only three hospitals, which may limit the generalizability of our findings. Although our genomic data suggested possible genetic determinants, alternative explanations, including heteroresistance, inoculum effects, regulatory changes outside the loci examined and variability in susceptibility testing, cannot be excluded. Furthermore, while our results suggest a potential association between the *psr-pbp5* locus and lower cephalosporin MICs, the observed variability indicates that additional genetic or regulatory factors may contribute to cephalosporin susceptibility. Functional studies will therefore be required to establish the underlying mechanisms and determine the specific contribution of PBP5. Finally, given the clinical relevance of *E. faecalis* and its predominantly ampicillin-susceptible phenotype, further studies are warranted to characterize the prevalence and determinants of low cephalosporin MICs in this species.

In conclusion, our study demonstrates that low cephalosporin MICs occur in a substantial proportion of ampicillin-susceptible *E. faecium* and *E. lactis* isolates from diverse phylogenetic backgrounds and independent hospital settings, indicating that this phenotype is not restricted to a particular clone or epidemiological context. These findings challenge the notion of uniformly high intrinsic cephalosporin resistance in enterococci and reveal previously underappreciated heterogeneity in β-lactam susceptibility. Our findings are exploratory and do not justify changes to routine clinical practice. Rather, they support standardized susceptibility testing in research settings and experimental validation of candidate mechanisms before clinical implications can be established. Given the use of combinations such as ampicillin plus ceftaroline or ampicillin plus ceftriaxone in some clinical settings, these observations warrant targeted preclinical and mechanistic studies to better define their activity against enterococci.

## ACKNOWLEDGEMENTS

We are grateful to the staff of the Genomics Unit at the Centre for Genomic Regulation (Barcelona, Spain) and Eurofins Genomics (Ebersberg, Germany) for sequencing. We also want to acknowledge the CERCA Program/Generalitat de Catalunya.

## SUPPLEMENTARY MATERIAL

**Data S1.** JSON-formatted file including all clinical metadata and whole-genome sequencing (WGS) analyses (MLST, antimicrobial resistance genes and virulence factors) for ampicillin-susceptible *E. faecium* and *E. lactis* isolates.

**Data S2.** JSON-formatted file including all clinical metadata and whole-genome sequencing (WGS) analyses (MLST, antimicrobial resistance genes and virulence factors) for ampicillin-resistant *E. faecium* and *E. lactis* isolates.

**Figure S1.** Heatmap showing the prevalence of antimicrobial resistance genes, point mutations and virulence factors in ampicillin-susceptible *E. faecium* isolates exhibiting low and high ceftriaxone MICs. The color scale represents prevalence percentage ranging from 0% (red) to 100% (green). No significant differences in genetic features were observed between both groups.

**Figure S2.** Unrooted phylogeny of the *pbp5-psr* locus of *E. faecium* and *E. lactis* isolates analyzed in this study. Branch support values represent the percentage of bootstrap replicates supporting each node and are shown only for branches with ≥ 75% support. Ampicillin susceptibility phenotype, ceftriaxone MIC and *psr* type are also indicated.

**Figure S3.** Graphical representation of the *psr-pbp5* platform deletion generated using clinker. Genes are represented by arrows indicating their orientation and homologous genes are connected between genomes of different isolates.

**Figure S4.** Individual and concatenated unrooted phylogenies of the amino acid sequences of CroS, CroR, MurAA, PBPA, PBPF, PONA, Stk and StpA. Where applicable, branch support values represent the percentage of bootstrap replicates supporting each node and are shown only for branches with ≥75% support. MLST, ampicillin susceptibility phenotype and ceftriaxone MIC are also indicated.

**Table S1.** Univariate analysis of clinical variables associated with ampicillin-susceptible isolates exhibiting low ceftriaxone MICs compared with those exhibiting high ceftriaxone MICs. Continuous variables were compared using the Mann-Whitney U test and categorical variables were assessed using Fisher’s exact test.

**Table S2.** Univariate analysis of genomic variables (ARGs and VFs) associated with ceftriaxone MIC in ampicillin-susceptible *E. faecium* isolates using Fisher’s exact test.

**Table S3.** Predicted proteins corresponding to genes previously associated with cephalosporin resistance. The table lists the query gene, the corresponding protein-coding sequence identifier in each isolate and the predicted amino acid sequence.

**Table S4.** Percent identity values between all PBP5 sequences identified in this study. Percent identity values are derived from pairwise alignments using the Needleman-Wunsch algorithm.

**Table S5.** Predicted *psr* and *pbp5* locus identified across genome assemblies by BLASTN searches. For each predicted operon, the table reports the assembly, contig, strand orientation, genomic coordinates, operon length and the nucleotide sequence identity, alignment coverage and hit length for each gene.

**Table S6.** Genes and functional annotations within the chromosomal deletion including the *pbp5* operon. The table shows the corresponding *E. faecium* SRR24 reference protein, OGS identifier, functional description, COG category, Pfam domains and VFs.

