## Supplementary material for "Widespread occurrence of ampicillin-susceptible *Enterococcus faecium* and *Enterococcus lactis* clinical isolates with low MICs to cephalosporins from Spain and Portugal": FigureS4.pdf

### CroR phylogeny

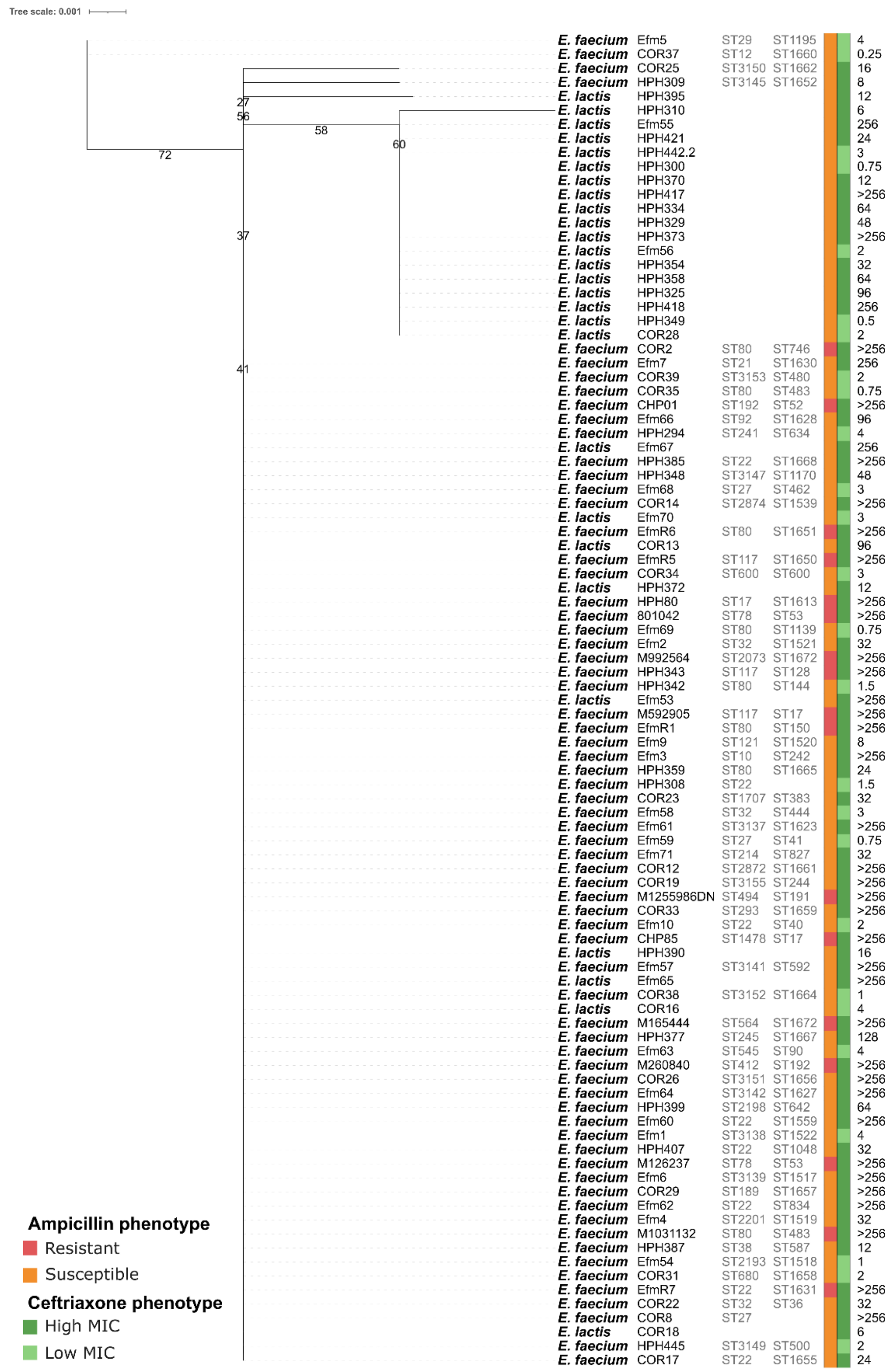

### CroS phylogeny

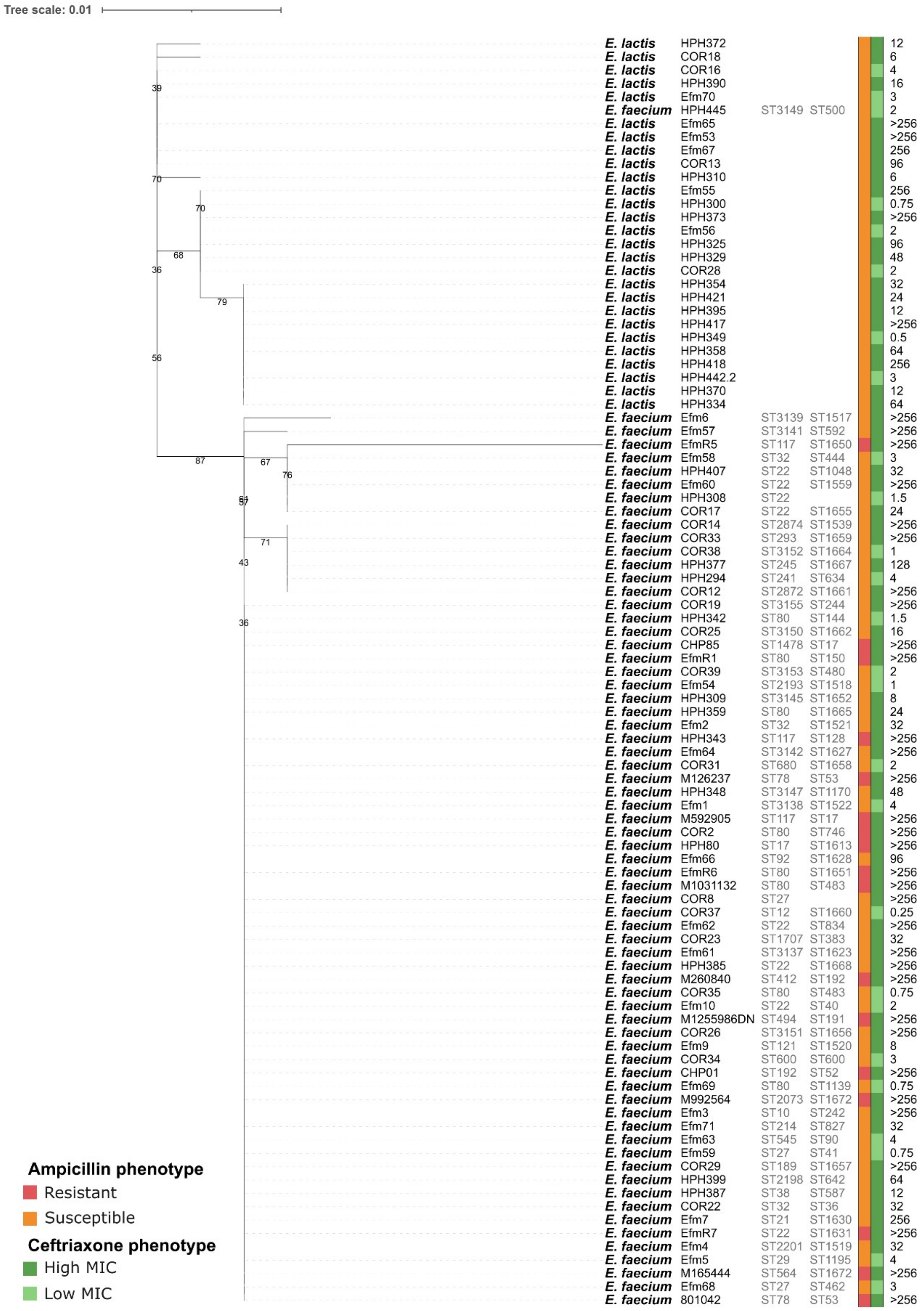

### MurAA phylogeny

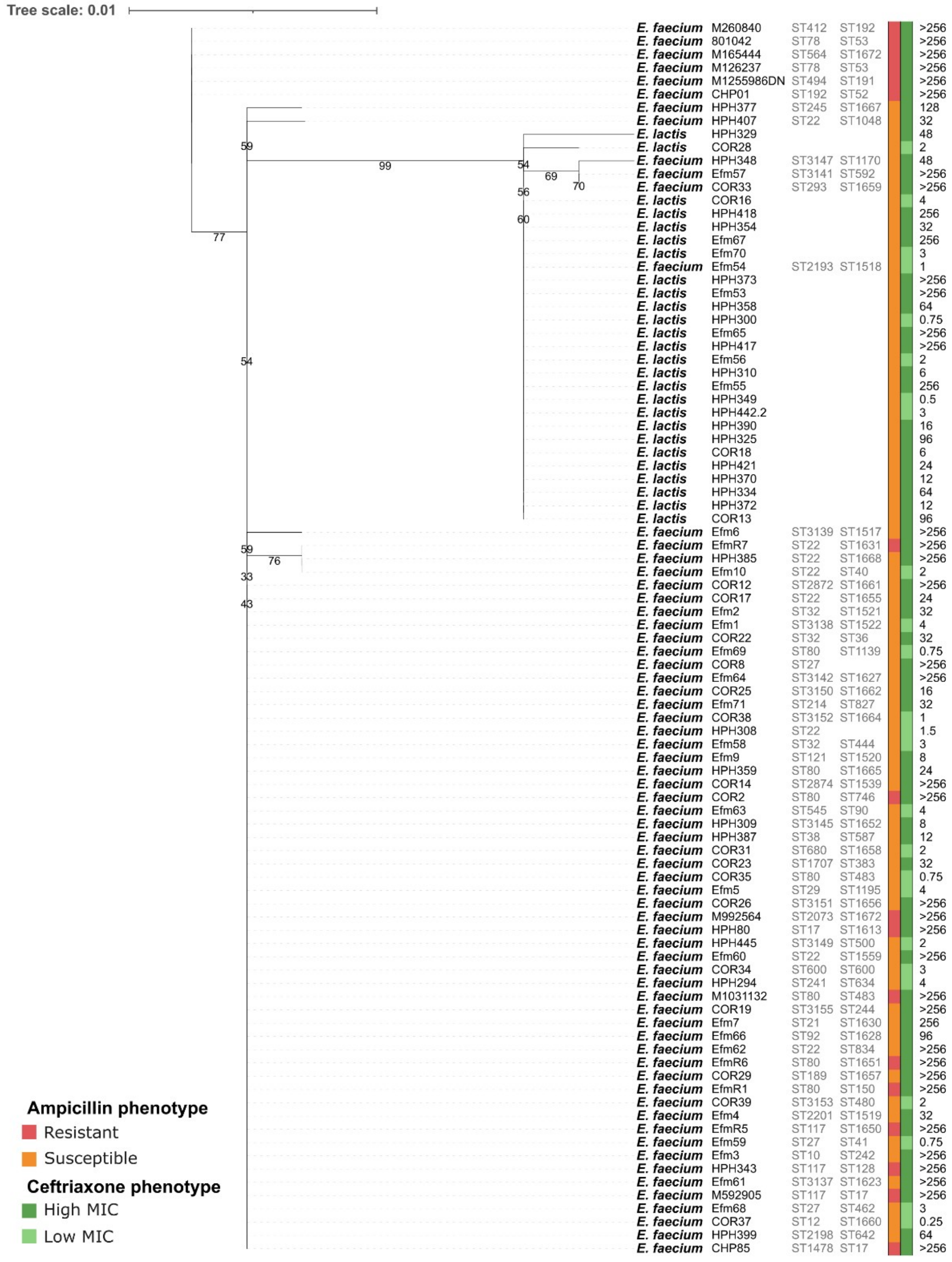

#### PBPA phylogeny

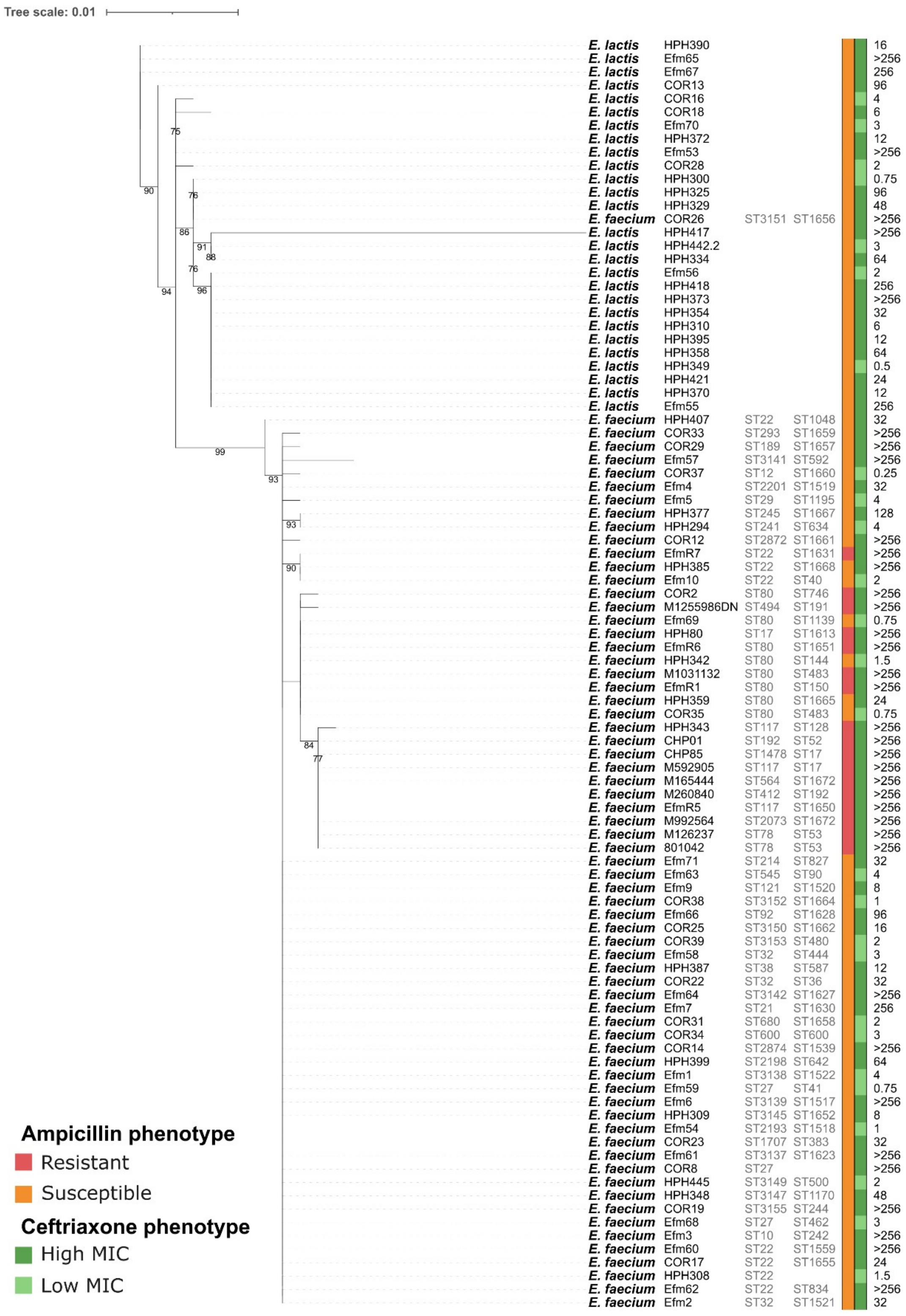

### PBPF phylogeny

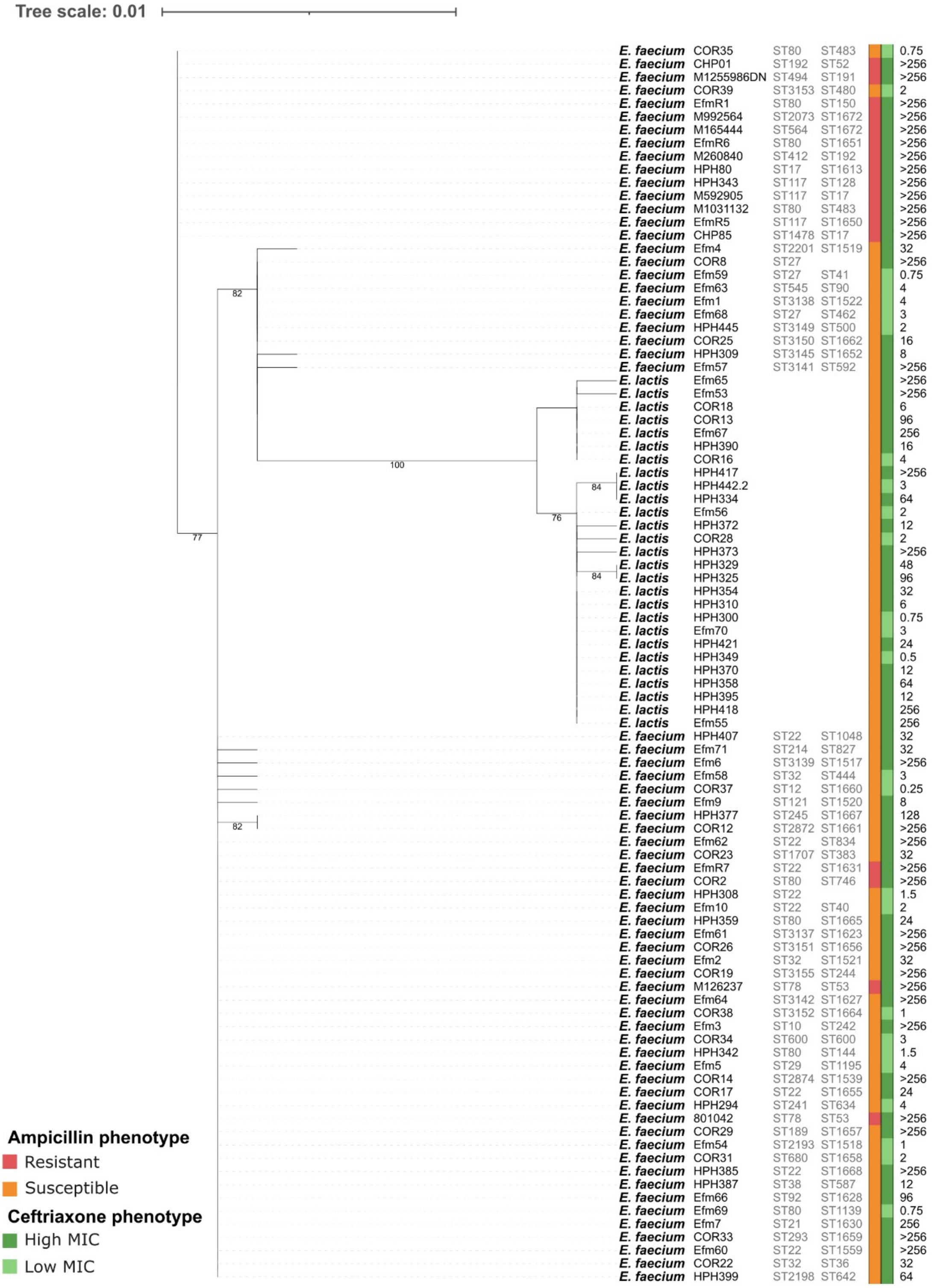

### PONA phylogeny

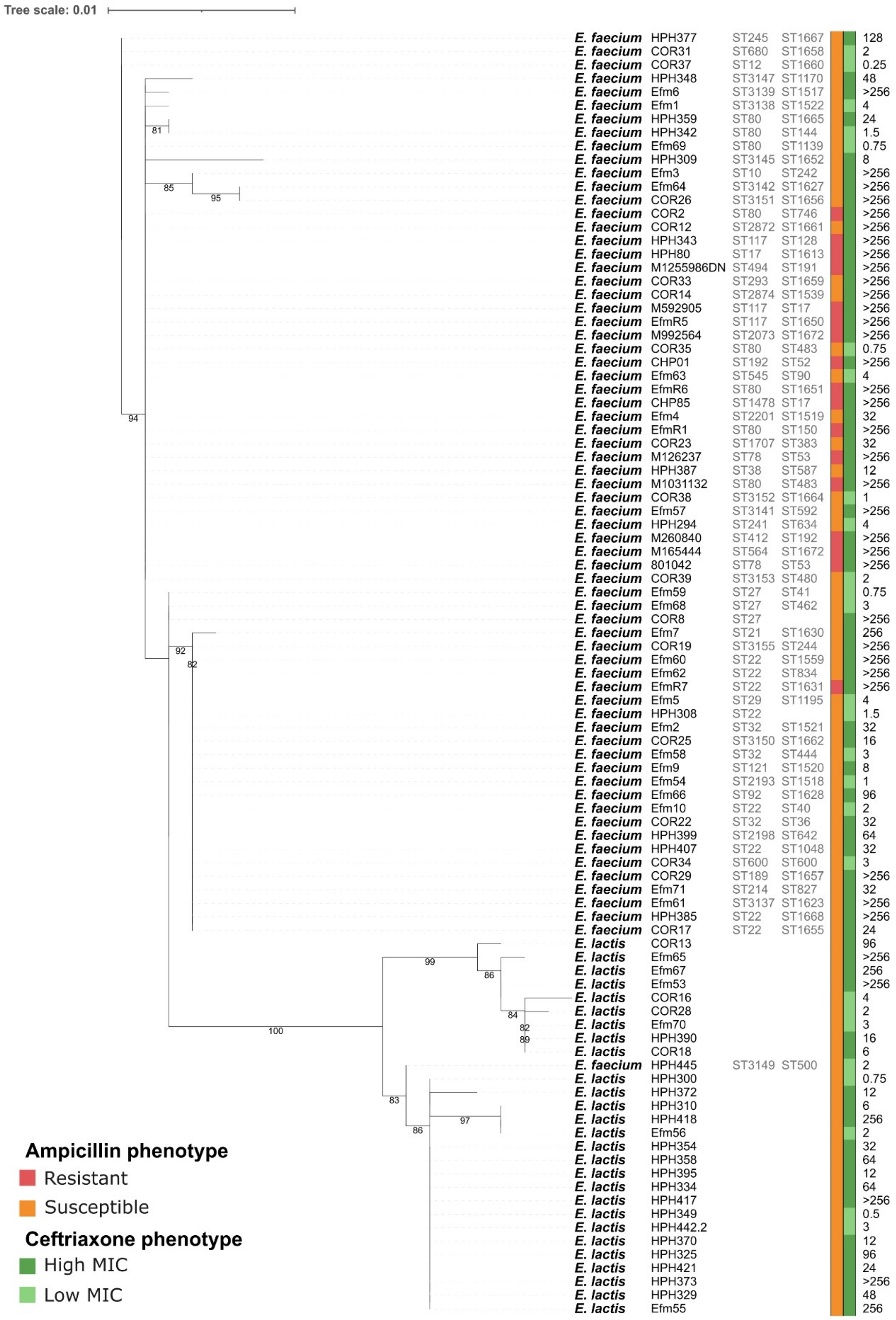

### Stk phylogeny

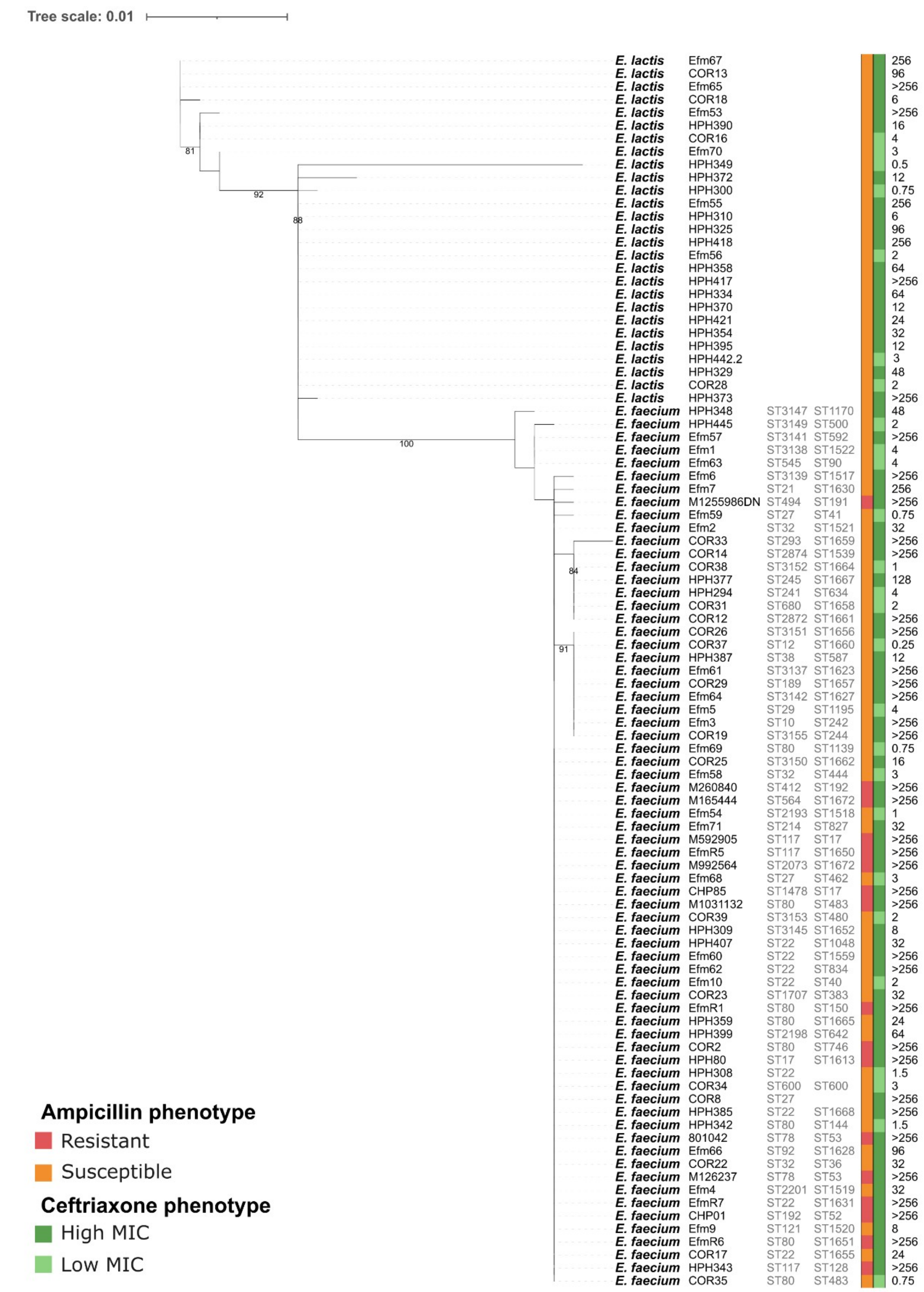

### StpA phylogeny

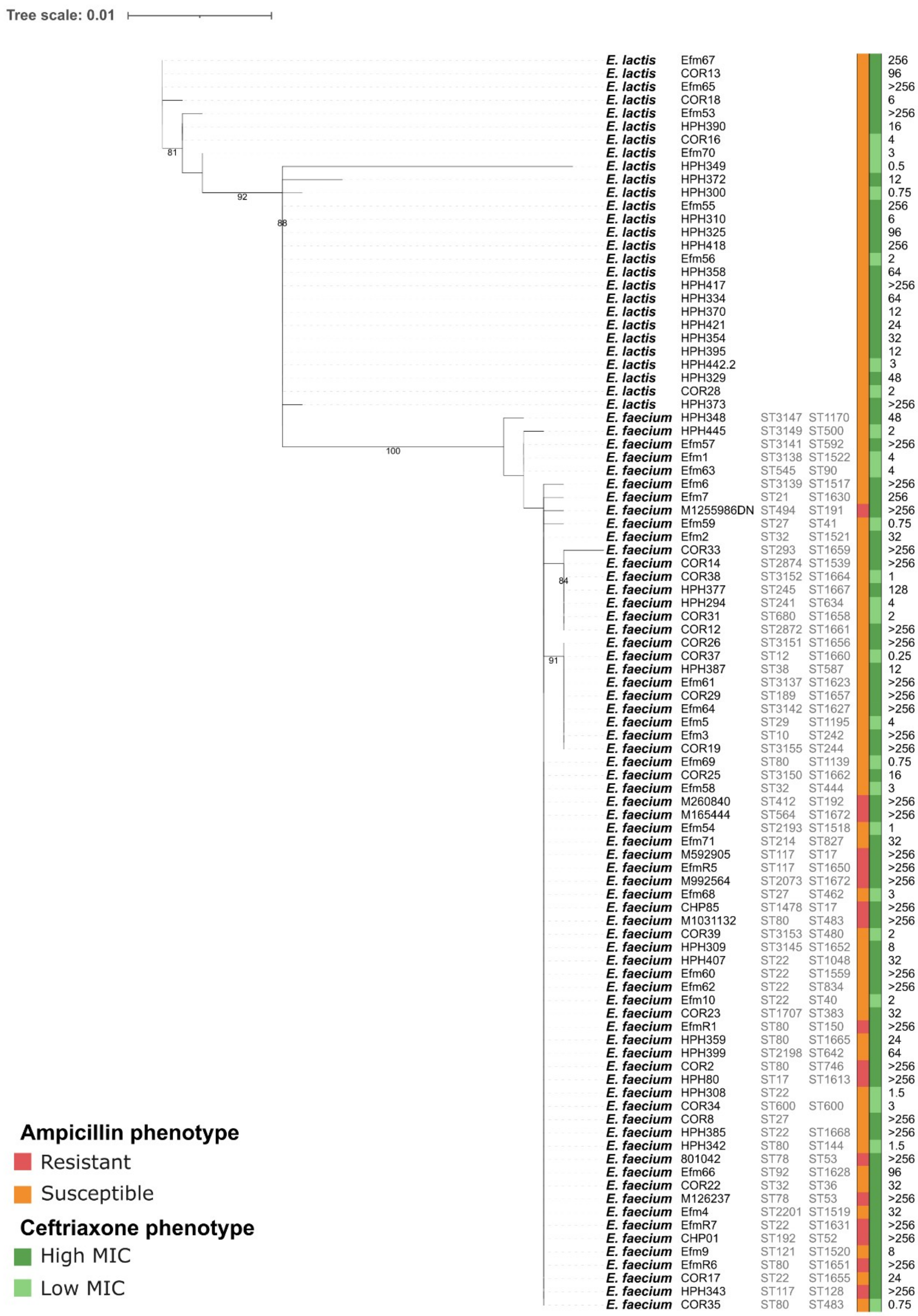

### Concatenated phylogeny

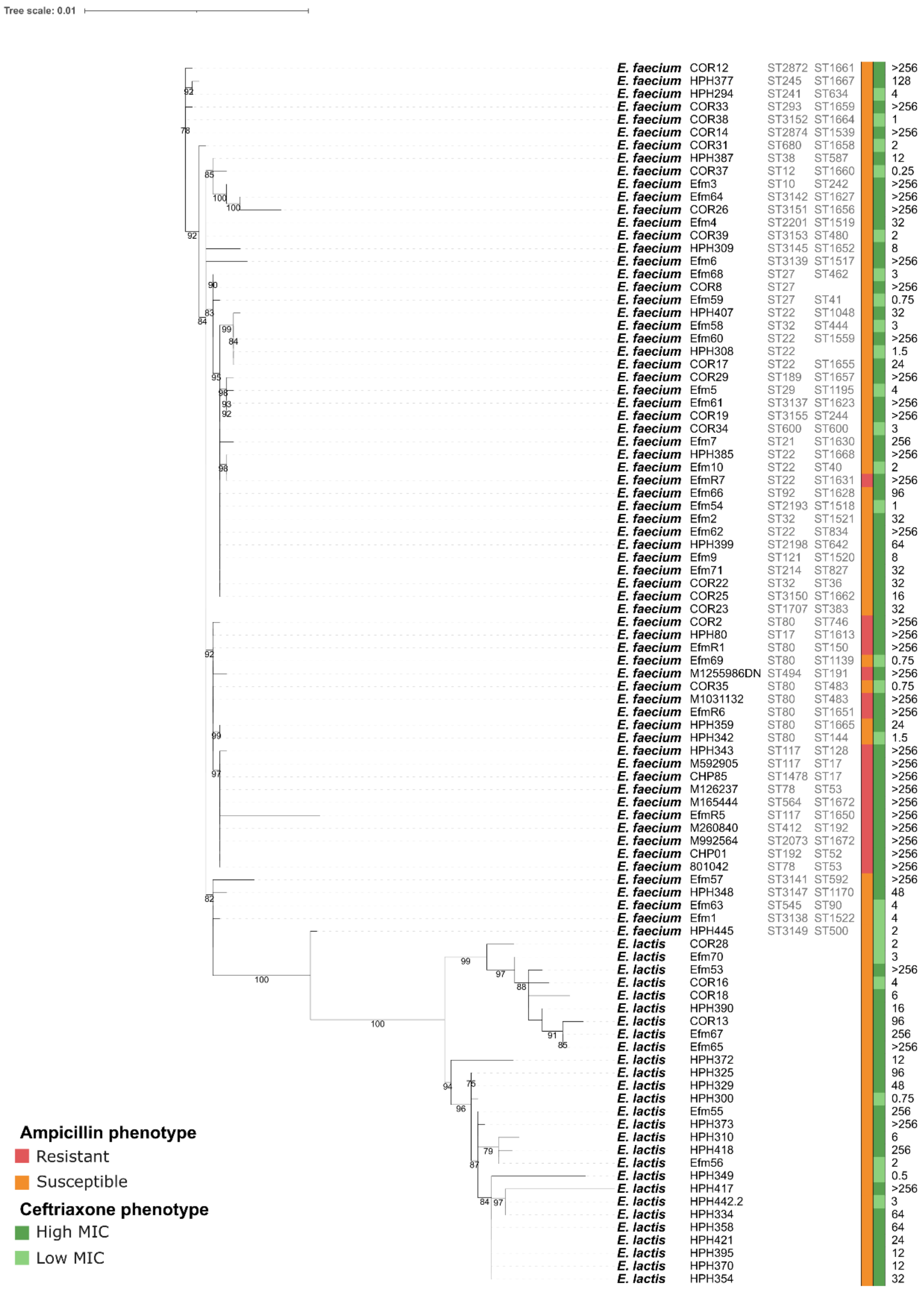
