## Supplementary figures and images for "Widespread occurrence of ampicillin-susceptible *Enterococcus faecium* and *Enterococcus lactis* clinical isolates with low MICs to cephalosporins from Spain and Portugal"

### FigureS1.jpg

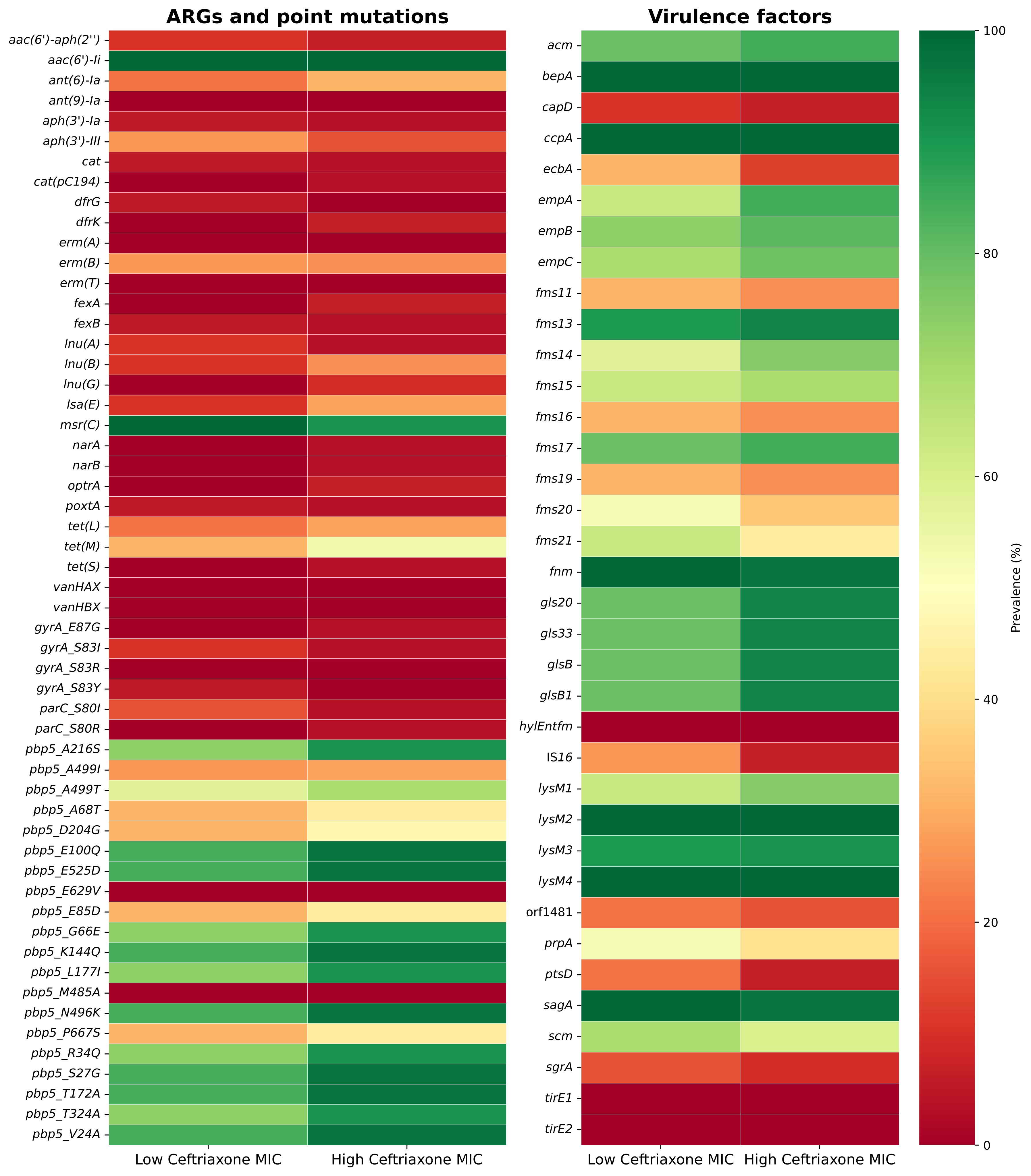

### FigureS2.jpg

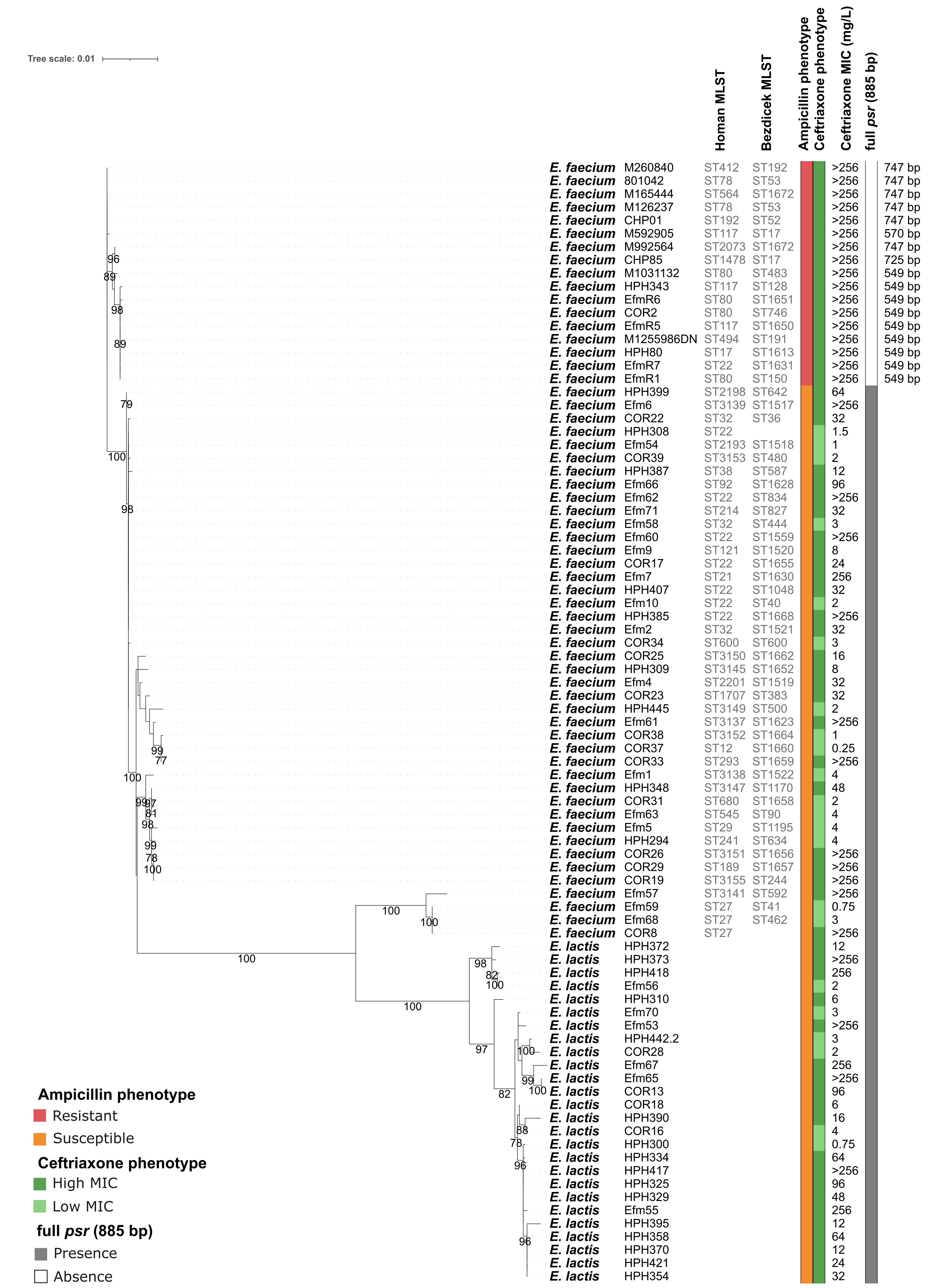

### FigureS3.jpg

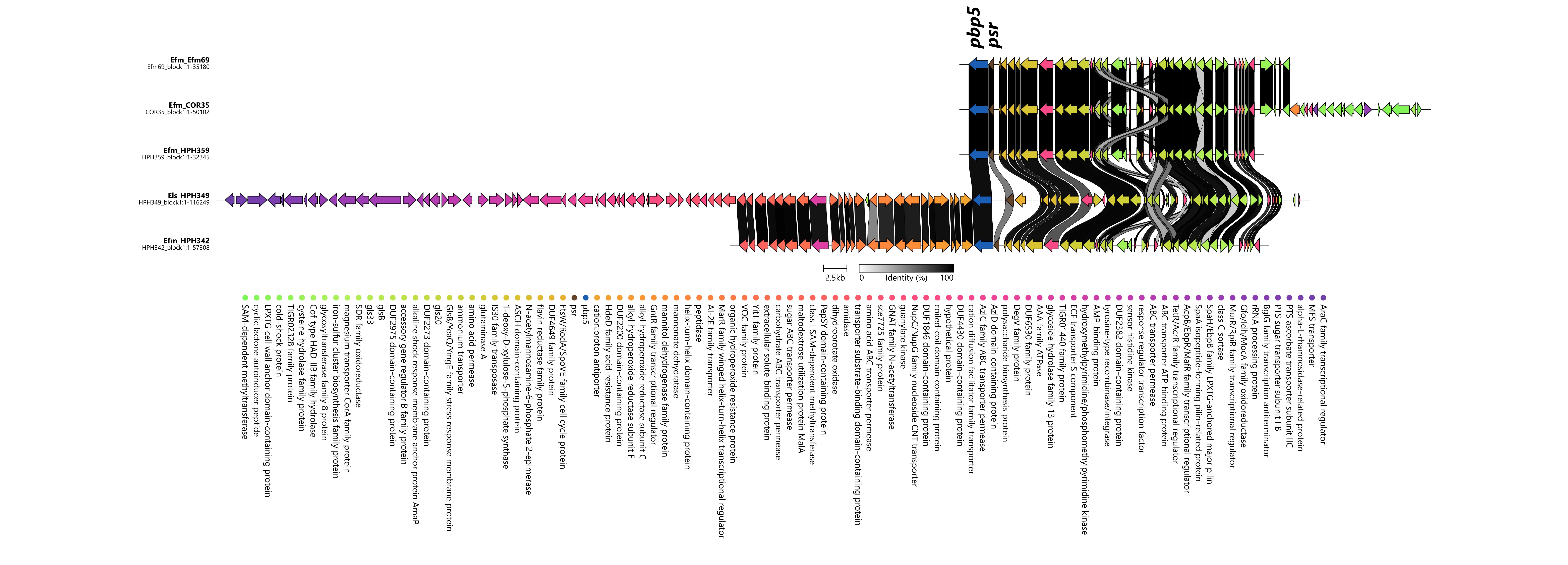
